# What makes clocks tick? Characterizing developmental dynamics of adult epigenetic clock sites

**DOI:** 10.1101/2024.03.12.584597

**Authors:** Rosa H. Mulder, Alexander Neumann, Janine F. Felix, Matthew Suderman, Charlotte A. M. Cecil

## Abstract

DNA methylation (DNAm) at specific sites can be used to calculate ‘epigenetic clocks’, which in adulthood are used as indicators of age(*ing*). However, little is known about how these clock sites ‘behave’ during development and what factors influence their variability in early life. This knowledge could be used to optimize healthy aging well before the onset of age-related conditions. Here, we leveraged results from two longitudinal population-based cohorts (*N*=5,019 samples from 2,348 individuals) to characterize trajectories of adult clock sites from birth to early adulthood. We find that clock sites (i) diverge widely in their developmental trajectories, often showing non-linear change over time; (ii) are substantially more likely than non-clock sites to vary between individuals already from birth, differences that are predictive of DNAm variation at later ages; and (iii) show enrichment for genetic and prenatal environmental exposures, supporting an early-origins perspective to epigenetic aging.

## Introduction

Individuals of the same age can show stark differences in health and age-related diseases, pointing to a distinction between chronological age and (biological) *ageing*. The reasons why people may age differently at a biological level are complex and multifactorial, e.g. reflecting the influence of genetic^1^, demographic (e.g. sex), lifestyle (e.g. smoking), and social (e.g. economic inequality) factors^2-4^. An increasingly popular method to capture age biologically is through the use of epigenetic data. DNA methylation (DNAm; the addition of methyl-groups to the DNA structure) is an epigenetic mechanism that regulates gene activity in response to both genetic and environmental influences, including prenatally^5-8^, and plays a key role in human development and health. Importantly, DNAm is highly age-dependent, and DNAm levels at specific sites can be used to calculate what are known as ‘epigenetic clocks’.

The first epigenetic clock was introduced by Horvath ^9^ a decade ago, and since then, several others have become available. These can be clustered according to the way they are built, or ‘trained’: while so-called ‘first-generation’ epigenetic clocks, including Horvath’s, were trained to predict chronological age^10-13^; ‘second-generation’ clocks (e.g. PhenoAge clock^14^, DNAmTL or ‘Telomere clock’^15^) were built to directly predict (indicators of) age-related diseases and/or mortality. Recently, the development of the DunedinPACE clock^16^ marks the beginning of ‘third-generation’ clocks, whereby longitudinal measurements of biomarkers of aging are incorporated in order to predict the ‘pace of aging’, such as triglycerides and urea nitrogen levels. Across the first- and second-generation clocks, it is possible to derive estimates of (i) epigenetic age – which tend to correlate with chronological age – as well as (ii) the extent to which epigenetic age deviates from chronological age, which can instead serve as a marker of biological ag*ing*. When epigenetic age is greater than chronological age, we call this epigenetic age acceleration, and when it is lower, we call it deceleration. Third-generation clocks provide an estimate of the rate of aging directly. Generally, findings using these clocks show that epigenetic age acceleration or increased rate of aging is associated with greater risk of age-related diseases and mortality in adulthood, whereas epigenetic age deceleration or decreased rate of ageing is associated with longevity and lower disease risk^17^. However, because the training and application of these epigenetic clocks has been almost exclusively confined to adult samples, their developmental roots remain largely unknown.

Relevant here is the fact that within clocks, tens or even hundreds of individual sites are combined to produce a single epigenetic age (or age acceleration) estimate. Yet, we know very little about how these individual sites ‘behave’ during development and whether they share common characteristics that set them apart from non-clock sites, in terms of: (i) their developmental trajectory; (ii) how early they begin to vary between individuals; and (iii) which factors may explain this variability. Given that the developmental characteristics of DNAm sites are related to their underlying biological function^18^, investigating clock sites from a developmental perspective could improve our understanding of ageing biology and, ultimately, how to optimize healthy ageing early on, well before the onset of age-related conditions.

To address these questions, we decomposed adult epigenetic clocks into their individual ‘components’ (i.e. sites), to characterize timing and patterns of change in their methylation levels over the first two decades of life, using results from two large-scale population-based cohorts with longitudinally assessed DNAm. We examine these developmental dynamics for commonly used first-generation and second/third-generation clocks. Further, we conduct enrichment analyses to test whether clock sites show different patterns of developmental variability compared to non-clock DNAm sites, and whether early variation in clock sites is linked to genetic and prenatal environmental influences, based on available multi-cohort epigenome-wide association studies (EWAS).

## Results

We describe here results for seven commonly used epigenetic clocks in humans, including first-generation clocks, i.e. Horvath’s clock^9^ (353 sites), Hannum’s clock^10^ (71 sites), Weidner’s clock^12^ (102 sites), and Zhang’s Elastic Net clock^11^ (514 sites), as well as second/third-generation clocks, i.e. PhenoAge^14^ (513 sites), the DNAmTL^15^ (140 sites), and DunedinPACE^16^ (173 sites). Since second- and third-generation clocks are both trained on (indicators of) age-related diseases or mortality, instead of age itself, these clocks were grouped together. In total, the first-generation clocks consist of 967 unique sites, and the second/third-generation clocks consist of 821 unique sites. For three developmental features (DNAm change, inter-individual differences, early origins) examined in this section, we report (i) the prevalence of these features for all DNAm sites measured across the genome as a benchmark (as described in a previous analysis^18^), and (ii) the prevalence for clock sites (separately for first generation and second/third generation sites), and whether this differs significantly from *non-clock* sites, based on enrichment analyses. An overview of findings is displayed in **Figure 1**, and enrichment results for individual clocks are provided in **Supplementary Table 1**.

**Figure 1.**
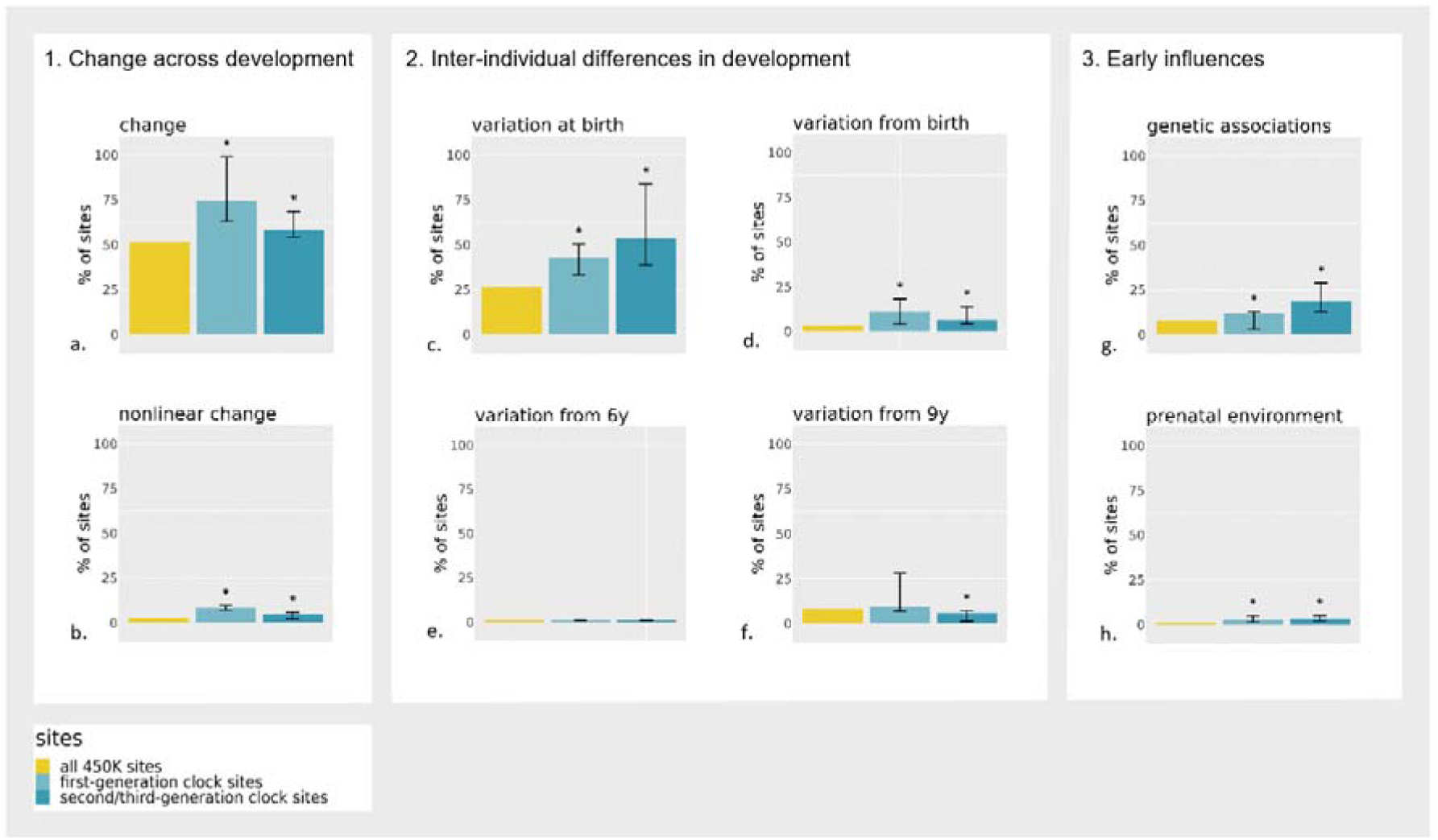
Overview of comparisons made between clock sites (first-generation and second/third-generation) versus non-clock sites, on the presence of a) DNAm change from birth to 18; b) nonlinear DNAm change at age 9; c) inter-individual DNAm differences at birth; d) inter-individual differences in DNAm change starting from birth; e) inter-individual differences in DNAm change starting from age 6 years (mid-childhood); f) inter-individual differences in DNAm change starting from 9 years (late childhood); g) genetic associations with DNAm at birth; and h) prenatal environmental association with DNAm at birth. The error bars indicate the range of percentages found across clocks; significant (*p*<0.05) differences between clock sites and non-clock sites on the array based on enrichment analyses are depicted with an asterisk (*).

### 1. Do clock sites change across development?

We used summary statistics from our EWAS of longitudinal DNAm across development to characterize epigenetic clock sites, based on repeatedly assessed DNAm from two population-based cohorts (the Generation R Study^19,20^ and the Avon Longitudinal Study of Parents and Children (ALSPAC)^21,22^). Analyses comprised 5,019 DNAm samples collected from 2,348 individuals at multiple time-points between birth and early adulthood^18^. DNAm was measured with the Illumina 450K array (>450,000 sites). As a first step, we characterize linear and non-linear dynamics of DNAm of first- and second/third-generation clock sites over the first two decades of life.

#### DNAm change during development

We considered a site as showing evidence of change during development if it associated with age from birth to early adulthood at a genome-wide significant level (Bonferroni-corrected threshold *p*<1×10^−07^; **Figure 2**). Based on this threshold, 52% of all DNAm sites measured on the array showed change during development, with DNAm increasing in 16% and decreasing in 36% of sites^18^. First-generation clocks were highly enriched with sites that change during development, especially with sites with increasing DNAm, compared to non-clock sites (75% [35% increasing; 40% decreasing], range across clocks=63-99%, OR=2.8 [95% CI=2.4-3.2], *p*=1.32×10^−49^). Second/third-generation clock sites were also enriched for (increasing) change during development (58% [30% increasing; 28% decreasing], range=54-68%, OR=1.3 [95% CI=1.1-1.5], *p*=8.90×10^−05^) (**Figure 1**). Overall, increasing sites tended to have positive coefficients in clock models and decreasing sites tended to have negative coefficients (i.e. contribute positively or negatively to the age(*ing*) estimate), which indicates that the direction of DNAm change in development typically seemed consistent with the direction of change in adulthood as estimated by epigenetic clocks (first-generation clock sites: OR=12.5, [95% CI=8.9-17.9], *p*=1.26×10^−58^; second/third-generation clock sites: OR=2.4 [95% CI=1.6-3.5], *p*=4.06×10^−06^; **Supplementary Figure 1**).

**Figure 2.**
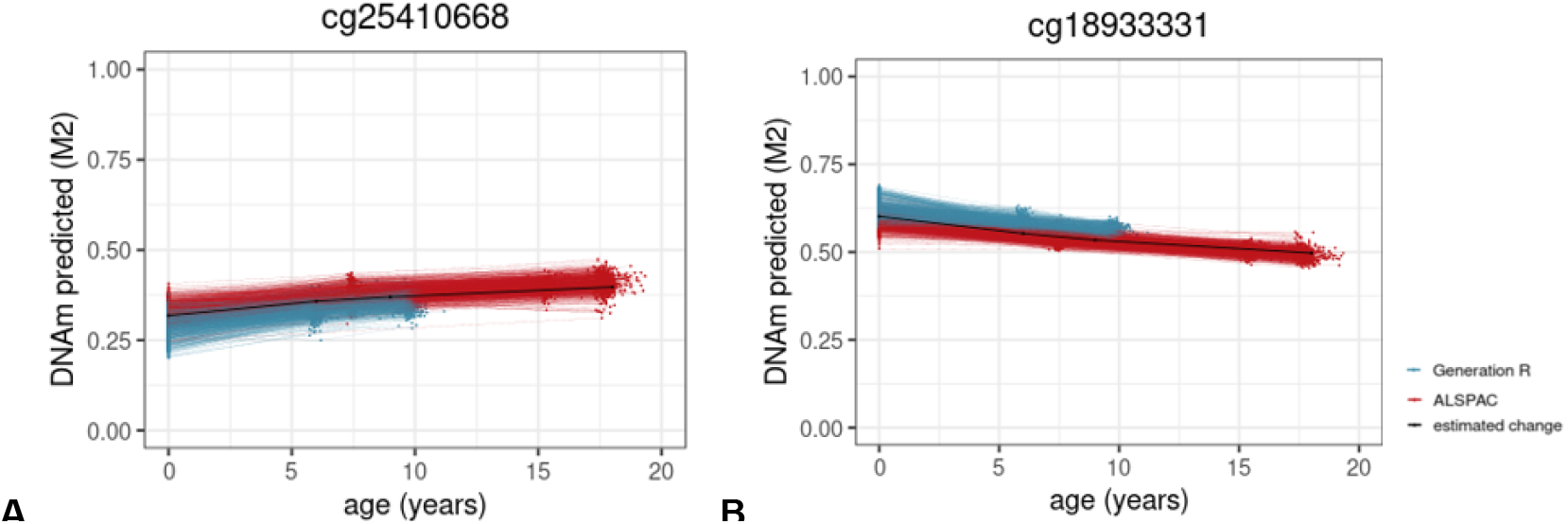
Examples of (**A**) a site from the Zhang’s and Hannum’s clock for which DNAm increases across development and (**B**) a site from Zhang’s clock for which DNAm decreases across development. Each line represents an individual’s predicted level of DNA methylation over time. The black line represents the group-level DNAm value over time, indicating that DNAm levels increase late childhood, and then decreases.

#### Non-linear change during development

Non-linear change was detected in a longitudinal mixed model that included slope changes at ages 6 (mid-childhood) and 9 years (late childhood) (*p*<1×10^−07^). Across all sites measured on the array, 11% of sites showed significant non-linear change during development^18^. In the majority of cases (8%), DNAm levels increased or decreased from birth to age 6, after which they remained stable. First- and second/third-generation clock sites were enriched for this type of non-linearity with 23% (range=14-43%, OR=3.6 [95% CI=3.0-4.1], *p*=3.22×10^−49^) and 18% (range=12-27%, OR=2.5 [95% CI=2.1-3.0], *p*=1.76×10^−19^) of sites, respectively. Of note, since DNAm was drawn from cord blood at birth and peripheral blood later on, this non-linearity may partly reflect tissue differences rather than developmental changes per se. Yet, a similar enrichment pattern was observed for sites showing non-linear change later in development, at age 9: this type of non-linear pattern was present in 3% of sites on the array (*p*<1×10^−07^), whereas it was 8% (range=7-10%) in first-generation clock sites, and in 5% (range=2-6%) in second/third generation clock sites – a significant enrichment in both groups compared to non-clock sites (OR=3.3 [95% CI=2.6-4.1], *p*=6.40×10^−18^ and OR=1.8 [95% CI=1.2-2.5], *p*=1.46×10^−03^, respectively; **Figure 1**). An example of a non-linear DNAm pattern is shown in **Figure 3**.

**Figure 3.**
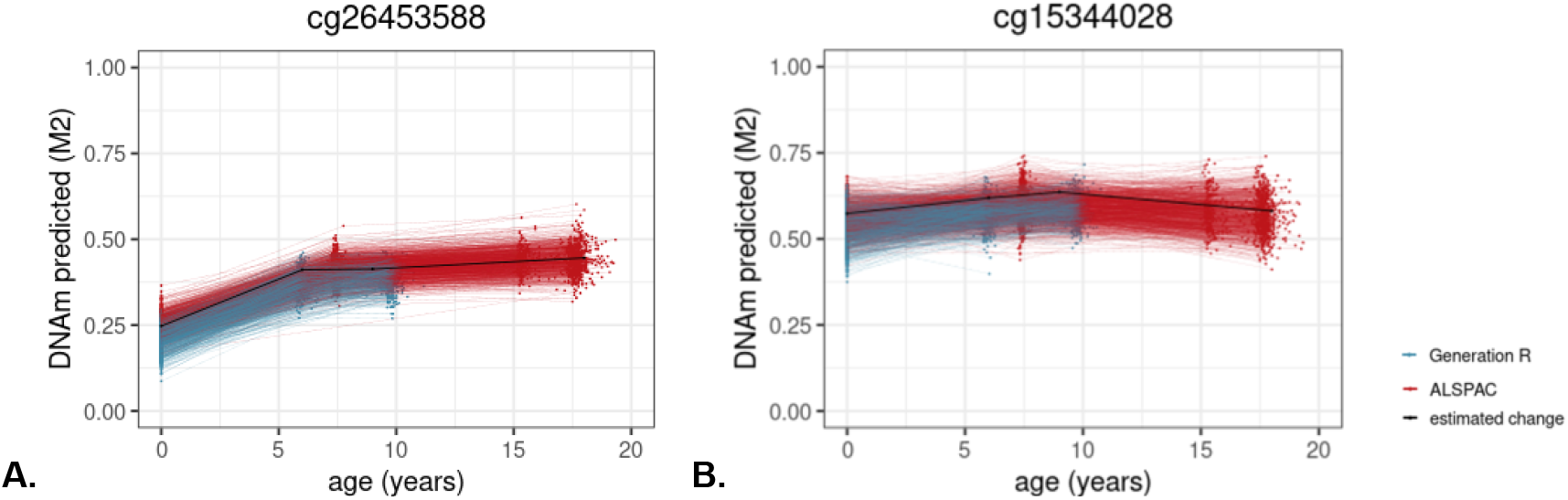
Example of a site from (**A**) Horvath’s clock, for which DNAm increases until age 6, after which no significant change happens up to age 18, and a site from the (**B**) the PhenoAge clock, for which DNAm increases until age 9, after which it decreases.

### 2. Do the developmental trajectories of clock sites differ between individuals?

We then asked how clock sites varied *between individuals* throughout development, namely: (i) to what extent clock sites already show inter-individual differences in DNAm levels at birth; and (ii) whether clock sites show inter-individual differences in the rate of DNAm change across development, and if so, at what age this begins to occur.

#### Inter-individual differences in DNAm levels at birth

Inter-individual differences at birth were quantified using a random intercept in the longitudinal mixed model (*p*<1×10^−07^). For all 450K sites, 23% of sites show significant individual differences at birth^18^. This proportion was 29% for first-generation clock sites (range=14-34%), and 47% of the second/third-generation clock sites (range=33-76%) – indicating an enrichment of these sites (OR=1.3 [95% CI=1.2-1.6], *p*=4.72×10^−05^; OR=2.9 [95% CI=2.6-3.4], *p*=8.46×10^−50^, respectively; **Figure 1**). An example of individual differences in DNAm at birth is shown in **Figure 4**.

**Figure 4.**
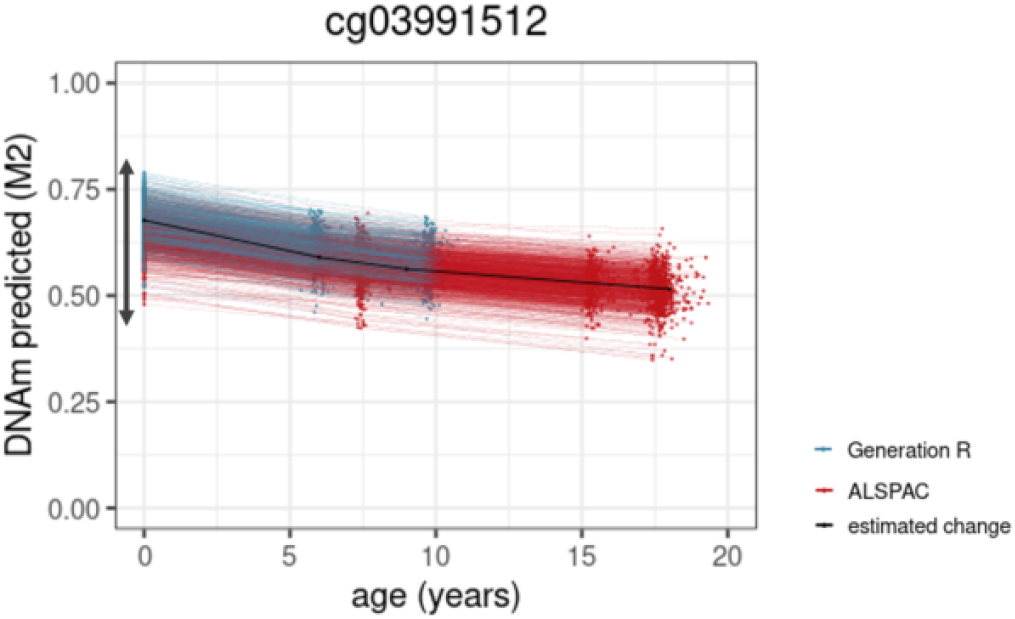
Example of a site from the PhenoAge clock. The arrow indicates the range of inter-individual differences in level of DNA methylation at birth.

#### Inter-individual differences in rate of DNAm change across development

We used the random slope estimates from the longitudinal mixed models to identify significant (*p*<1×10^−07^)inter-individual differences in rate of change at birth, mid-childhood (age 6), and late childhood (age 9) (**Figure 1**)^18^.

i. *Inter-individual differences beginning at birth*: Only around 3% of all 450K sites show inter-individual differences in rate of change from birth onward^18^. This proportion was significantly higher in both first-generation (11% of sites [range=4-18%]; OR=3.6 [95% CI=2.9-4.4], *p*=5.27×10^−26^) and second/third-generation clocks (7% of sites [range=4-14%]; OR=2.0 [95% CI=1.5-2.7], *p*=3.54×10^−06^). An example of this pattern can be found in **Figure 5A**.
ii. *Inter-individual differences beginning in mid-childhood:* Sites with inter-individual differences in rate of change starting in mid-childhood were sparse, comprising 0.2% of all 450K sites^18^. In first- and second/third-generation clocks, these numbers were similarly low, with 0.1% (range=0-0.2%, OR=0.6 [95% CI=0.0-3.5], *p*=1.00), and 0.4% (range=0-0.4%, OR=2.2 [95% CI=0.4-6.4], *p*=0.16) of sites, respectively (**Figure 5B**).
iii. *Inter-individual differences beginning in late childhood:* Last, 8% of all 450K sites showed inter-individual differences in rate of change starting in late childhood^18^. In first-generation clocks, these numbers were comparable (9% of sites (range=7-28%), OR=1.1 [95% CI=0.9-1.4], *p*=0.24), whereas they were somewhat lower (i.e. depleted) for second/third-generation clocks (6% [range=1-7%]; OR=0.7 [95% CI=0.05-0.9], *p*=0.02, **Figure 5C**). Taken together, these results indicate that clock sites show strong enrichment for inter-individual variability in rate of change starting from birth, but not at later time points in development.

**Figure 5.**
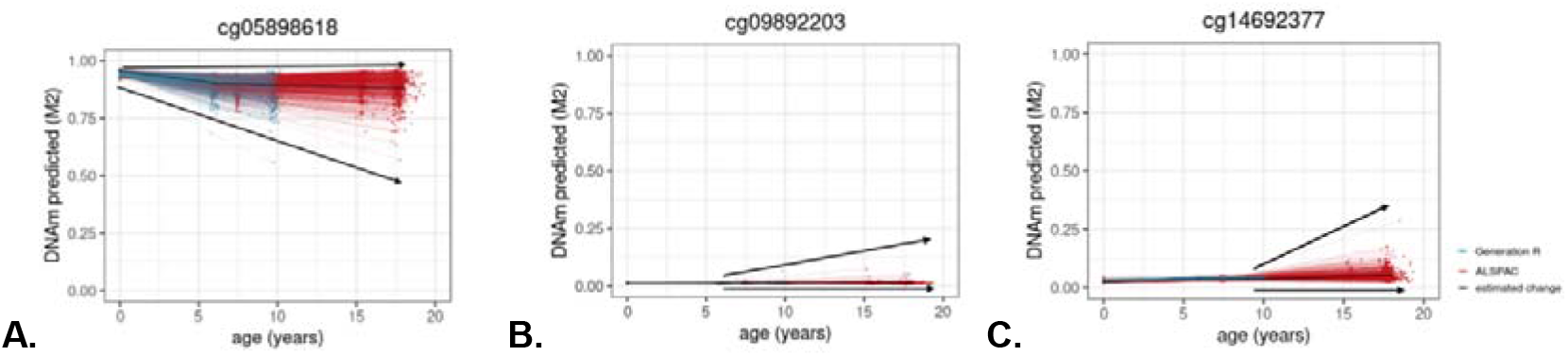
Examples of a site from Zhang’s (**A**) and Hannum’s (**B**) and the PhenoAge (**C**) clock, for which inter-individual differences in DNA methylation start to appear from birth (**A**), mid-childhood (**B**), and late childhood (**C**) onwards. The arrows indicate the range of inter-individual differences in rate of change.

#### Can DNAm levels at birth be used to predict DNAm levels in emerging adulthood?

For each site, we tested whether inter-individual differences in DNAm levels at birth were associated with inter-individual differences in rate of DNAm change up to early adulthood. Among clock sites that had any inter-individual differences at birth and in change, we found that almost all (97%) showed significant (*p*<1×10^−07^) correlations between the random intercept (i.e. individual differences in DNAm at birth) and slope (i.e. individual differences in rate of DNAm change), with absolute mean *r*=0.42 in first-generation clock sites and *r*=0.39 in second-generation in clock sites.

To illustrate the degree to which these correlations can be leveraged to predict DNAm inter-individual differences in early adulthood based on ‘baseline’ differences at birth, we refer to **Figure 5A**. Here, we can see that inter-individual differences at birth are small and not significant; yet, individuals who show higher (rank) DNAm levels at birth seem to remain high over time, whereas those with lower DNAm levels at birth show a more pronounced decrease in DNAm levels, resulting in exacerbated DNAm differences at age 18. Indeed, the correlation between inter-individual DNAm differences at birth and in DNAm change in this case was *r*=0.62, thus 39% of variation in DNAm at age 18 could already be predicted based on DNAm at birth.

### 3. Probing potential early influences on inter-individual differences in clock sites

To better understand what potential factors may drive DNAm variability at birth in clock sites, we examined enrichment for genetic and prenatal environmental factors.

#### Genetic influences

We tested whether clock sites are associated with known methylation quantitative trait loci at birth (meQTL; i.e. common genetic variants associated with DNAm variation), based on genome-wide association analyses of DNA methylation at birth in the ALSPAC cohort^23^. On an epigenome-wide level, 8% of sites have been linked to an meQTL at birth in this database. These percentages were higher in first-generation clock sites (12% [range=3-13%]; OR=1.6 [95% CI=1.2-1.9], *p*=1.59×10^−05^) and second/third-generation clock sites (19% [range=13-29%]; OR=2.7 [95% CI=2.2-3.2], *p*=5.81×10^−23^) (**Figure 1**). As a sensitivity analysis, we repeated this analysis on an meQTL set^8^ that was based on DNAm at all ages, resulting in many more identified meQTLs (31% on an epigenome-wide level), which similarly showed an enrichment of meQTL associations amongst first-(62% [range=50-71%], OR=3.8 [95% CI=3.3-4.4], *p*=1.96×10^−93^) and second/third-generation (51% [range=48-56%], OR=2.4 [95% CI=2.1-2.7], *p*=1.63×10^−34^) clock sites. Two examples of genetic influences are provided in **Figure 6**.

**Figure 6.**
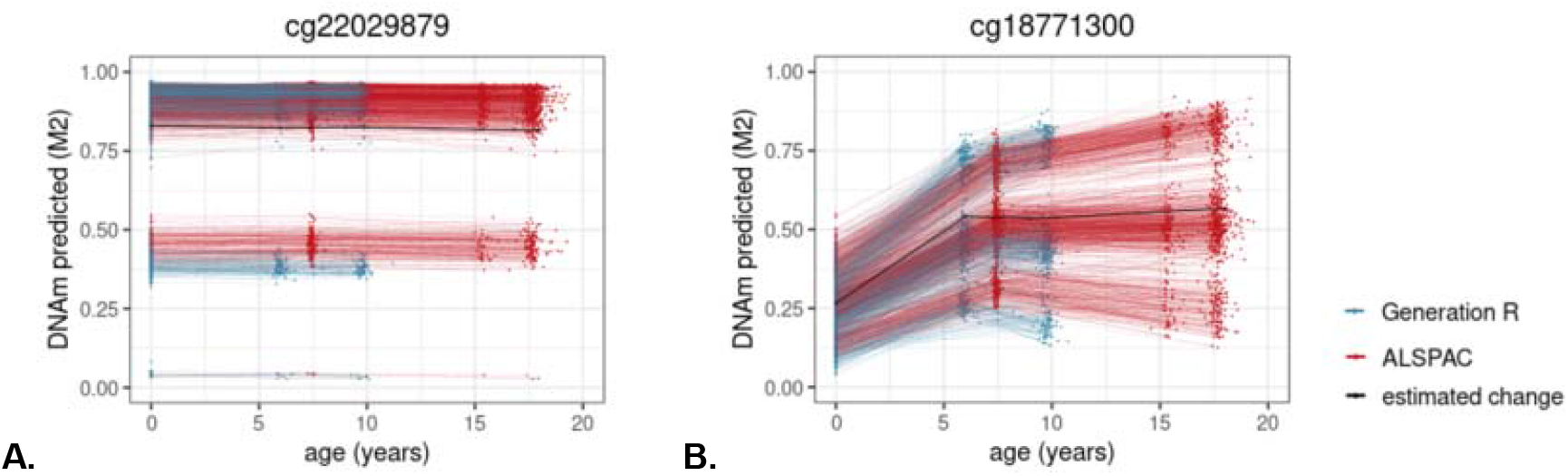
An example of a site from the DunedinPACE clock (**A**) and the PhenoAge clock (**B**) showing individual DNAm differences at birth related to a genetic polymorphism, which appear maintained (**A**) or exacerbated (**B**) across development.

#### Prenatal exposures

To test whether variation in DNAm at clock sites may be sensitive to prenatal environmental factors, we combined findings from existing EWAS meta-analyses from the Pregnancy and Childhood Epigenetics (PACE) consortium^24^. Findings were included based on availability in the EWAS Catalog^25^. Prenatal exposures included sustained maternal smoking (*n*=2,620 associated sites)^26^, hypertensive disorders (*n*=920)^27^, overweight/obesity (*n*=159)^28^, body mass index (BMI; *n*=104)^28^, (pregnancy-related) anxiety (*n*=57)^29^, haemoglobin levels (*n*=40)^30^, and air pollution exposure (NO_2_, *n*=24 PM_2.5_, *n*=6)^31,32^. Altogether,1% of the sites measured on the 450k array have been related to a prenatal exposure. This proportion was significantly larger in first-generation clock sites with 3% (range=1-5%, OR=3.9 [95% CI=2.6-5.7], *p*=9.74×10^−10^) and in second/third-generation sites with 2% (range=1-5%, OR=2.8 [95% CI=1.6-4.4], *p*=1.77×10^−04^) (**Figure 1**).

## Discussion

We selected epigenetic clocks that have been robustly associated with age-related outcomes in adulthood, and ‘deconstructed’ them into their individual components (i.e. sites) to characterize the developmental trajectories of clock sites. To do so, we leveraged longitudinal data from over 2,300 individuals from the general population with repeated measurements of DNAm spanning the first two decades of life. We highlight here three key findings: (i) *Hidden heterogeneity*: we show that epigenetic clocks contain sites that diverge widely in their developmental trajectories, often changing in non-linear manner over time (i.e. changing at different rates for different developmental periods). This heterogeneity is consequential as it not only illustrates the developmental complexity of clock sites, but may also violate the modeling assumptions of epigenetic clocks. (ii) *Early variability*: we find that over a third of all clock sites exhibit inter-individual differences at the earliest stages of life, from birth – a significantly greater proportion than what is observed at non-clock sites across the genome – and that ‘baseline’ DNAm differences at birth are predictive of differences in DNAm *change* across development. This raises important questions about when and why differences between individuals in aging-relevant sites begin to emerge. (iii) *Genetic and prenatal influences*: clock sites do not only differ in their developmental trajectories, but also in the extent to which they relate to genetic and prenatal environmental factors. Amongst clock sites, genetic influences are prominent, but we also identify enrichment for prenatal environmental exposures including smoking, and maternal health, further supporting an early-origins perspective to epigenetic ageing.

Epigenetic clocks are typically used as a unitary construct (i.e. single score) to estimate chronological age or markers of biological ageing. Here, we show that underlying these clocks are sites that differ widely in their developmental patterns, both in terms of how they vary *over time* and *between individuals*. Despite this heterogeneity, we identify features that are enriched in clock sites compared to non-clock sites across the genome. As might be expected (given that these clock sites are selected to estimate age), clock sites were more likely to show age-related change, even in early life. Less expected was the finding that clock sites are also more likely than non-clock sites to change in a non-linear manner (i.e. at different rates during different developmental periods. This raises the question of whether the non-linearity of these sites is confined to development (with DNAm levels becoming more linearly associated with age in adulthood) or whether these sites continue to show non-linear patterns across the lifespan, and may not be picked up in the development of epigenetic clocks, which often rely on methods where linear associations are assumed. The existence of non-linear patterns across development may point to sensitive time-windows for age-related epigenetic processes and serve to highlight the complex developmental dynamics hidden within epigenetic clocks.

In addition to characterizing the way clock sites change across development, we also examined the extent to which they vary *between individuals* from birth to emerging adulthood. Unexpectedly, we see that clock sites show strong enrichment for inter-individual differences starting at or starting from birth, but not for inter-individual differences starting at later time points. In other words, clock sites were more likely than non-clock sites to show inter-individual differences both in (i) baseline levels of DNAm at birth, as well as (ii) the rate of change in DNAm from birth onward; in contrast, no enrichment was found for individual differences emerging in mid-or late-childhood. Second/third generation clocks showed the highest proportion of birth-varying sites (47%, compared to 28% in first-generation sites and 23% across the genome), which is surprising given that they are trained on aging-associated phenotypes in adults (e.g. age-related disease and mortality). Further, we found that amongst clock sites, an individual’s baseline levels of DNAm at birth was strongly predictive of their rate of DNAm change across development. Even when a clock site showed relatively little inter-individual differences at birth, based on a person’s rank it was still possible to predict their degree of DNAm change over time. These findings raise questions about which factors may drive early individual differences in clock sites in the first place.

We addressed this question by leveraging published data from large-scale multi-cohort epidemiological studies. We find most support for genetic influences on clock sites, consistent with previous research showing that epigenetic clocks are partly under genetic control^33^. In future, functional follow-ups of genetic variants associated with these early-varying clock sites may provide insights in the genetic processes related to epigenetic aging. We also observe an enrichment for prenatal exposures including maternal smoking and health. This is in line with findings from an earlier study that examined prenatal influences on age acceleration according to Horvath’s clock^34^. There, it was reported that maternal prenatal smoking associated with epigenetic age acceleration at birth, whereas prenatal maternal weight and BMI associated with epigenetic age acceleration in childhood. However, these associations were small in magnitude and could not be replicated in an independent cohort. This may be because, despite showing an overall enrichment for prenatal influences, clock sites differ from one another in the extent to which they associate with different exposures - information that may be obscured when using combined clock estimates. Future studies are needed to establish the role of the prenatal exposome on biological aging in adulthood.

Our findings should be interpreted in the context of a number of limitations. The first is that while epigenetic clocks relate to ageing, they are developed as predictive biomarkers and not based on causal relationships – hence we cannot ascertain that developmental DNAm trajectories of clock sites are causally related to later ageing. However, just as epigenetic clocks are valuable biomarkers of ageing, the developmental trajectories of clock sites may mark developmental processes that are relevant to ageing. Secondly, some non-clock sites may be ageing-related, but excluded during feature selection for containing redundant information with selected DNAm sites in epigenetic clocks. Thus, the reported comparisons may underestimate differences between ageing and non-ageing related DNAm sites. Thirdly, within epigenetic clocks, association estimates for each DNAm site are mutually adjusted for the effects of all other clock sites. In contrast, longitudinal change estimates from birth to childhood stem from separate regression models for each site. While this limits comparisons, we find that estimates were overall consistent with one another: DNAm levels which were found to increase between birth and early adulthood, also were more likely to show a positive association with age in epigenetic clocks, and vice versa. Fourthly, with the current enrichment approach we cannot establish if the identified genetic variants and environmental prenatal factors are causally associated with ageing; however, they do provide new testable hypotheses for research examining the early determinants of age-related methylation patterns. In future, it will be of interest to integrate data on genetic and environmental influences to test their joint and potentially interactive effect on clock sites. Furthermore, the inclusion of individual outcome data could allow us in the future to establish whether developmental variability in clock sites is associated with child and adolescent health-related outcomes, and how early these associations begin to manifest. A small number of studies investigating the relationship between epigenetic clocks and health outcomes in early life have produced inconsistent results^35-37^. It is possible that these inconsistencies, based on the use of epigenetic clock estimates, are due to the underlying heterogeneous relationships of individual clock sites with age-related exposures and outcomes, and could potentially be clarified by taking the site-level approach described here. Last, while the availability of repeated DNAm measures over two decades in the current study is unique, more extended data and/or combinations with other datasets will be necessary to better understand how development and aging across the lifespan are related to epigenetic variability.

In brief, we characterized for the first time how sites included in commonly used epigenetic clocks vary over the first two decades of life. We find that clock sites differ widely from one another in their developmental trajectory. They are, however, more likely than non-clock sites to show individual differences already from birth. Our enrichment analyses suggest that these early differences may largely be explained by genetic factors, and to a lesser extent, prenatal environmental influences, supporting an early-origins perspective of epigenetic aging.

## Methods

### Setting

Epigenetic data were obtained from two prospective population-based cohorts: the Generation R Study^19,20^ (Generation R) in The Netherlands and the Avon Longitudinal Study of Parents and Children^21,22^ (ALSPAC) in the United Kingdom. For a full description of the studies and sample selection we refer to the Supplementary Materials. In short, in Generation R, epigenetic data was available for 1,399 children, including 2,333 measurements at birth and/or 6 years and/or 10 years of age. The sample consisted of participants with parents born in the Netherlands. In ALSPAC, epigenetic data was available for 949 children, including 2,686 measurements at birth and/or 7 years and/or 17 years. All children had European ancestry. Together, these data formed a single dataset consisting of 2,348 children with 5,019 measurements.

### DNA methylation

DNA methylation was extracted from cord blood at birth in both cohorts, and from peripheral blood at mean ages 6.0 (SD=0.5) and 9.8 (SD=0.3) years in Generation R and at mean ages 7.5 (SD=0.2) and 17.1 (SD=1.0) years in ALSPAC. The EZ-96 DNAm kit (shallow) (Zymo Research Corporation, Irvine, CA) was used for bisulfite conversion on the extracted DNA. Samples were then processed with the Illumina Infinium HumanMethylation450 BeadChip (Illumina Inc., San Diego, CA). Quality control occurred separately in each cohort^18^ and data from both cohorts were then normalized together as a single set to minimize technical variability. Functional normalization was applied with the *meffil* package^38^. Analyses were restricted to 473,864 autosomal DNAm sites. DNAm levels were operationalized as beta values, ranging between 0 and 1, representing the ratio of methylated signal relative to the sum of methylated and unmethylated signal per CpG.

### Longitudinal models

For these analyses, we used the site-level summary statistics of two models. First, to study *overall change in DNAm throughout development* and *inter-individual differences at birth*, we used a linear mixed model (Model 1), defined as follows:

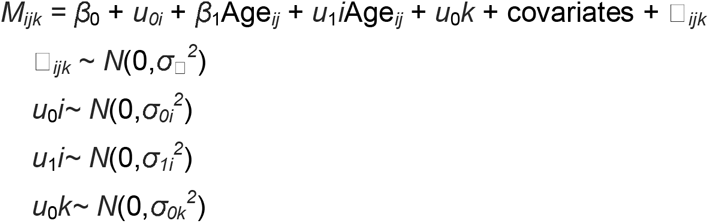

With participants denoted by *i*, time points by *j*, and sample plates by *k*. DNAm level is denoted by *M*, fixed intercept by *β*_0_, random intercept by *u*_0_*i*, fixed age coefficient by *β*_1_, random age coefficient by *u*_1_*i*, and last, random intercept for sample plate by *u*_0_*k*. The fixed age coefficient (*β*_1_) was used to estimate overall change in DNAm and the random intercept at the individual level (*u*_0_*i*) was used to estimate inter-individual differences in DNAm level at birth.

Second, to study *overall non-linear* change and *inter-individual differences in rate of change* at specific time-points, we used a linear mixed model including slope changes, to estimate non-linear DNAm trajectories (Model 2). Model 2 was defined as follows:

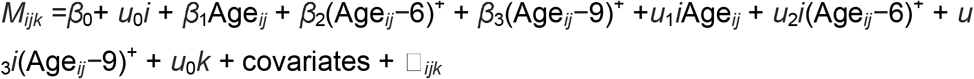

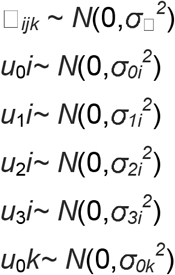

Here, a^+^□= □a if a□> □0 and otherwise 0. As such, β_1_ denotes the average change in DNAm per year from birth onwards, β_2_ represents the average change in DNAm per year from 6 years onwards after accounting for β_1_, i.e. the slope change or rate of change at age 6, and similarly β_3_ represents the average change in DNAm per year from 9 years onwards, after accounting for β_2_. With random effects *u*_1_*i, u*_2_*i*, and *u*_3_*i* inter-individual variation in slope (change) is denoted.

Analyses were performed using maximum likelihood estimation with the *lme4* package^39^ in R^40^. The models were fit with a diagonal random effect matrix to avoid convergence issues. *P-*values for fixed effects were obtained with *z-*tests of T-values; *p*-values for random effects were obtained by refitting the model without the random effect and comparing fit estimates in a likelihood ratio test. Significance thresholds were adjusted for the number of DNAm sites to *p*<1×10^−07^. Correlations between inter-individual differences at birth and in rate of change were obtained by applying Pearson correlations to the extracted random intercept best linear prediction (BLUP, or conditional modes) and random slope BLUP. All models were adjusted for batch (sample plate number), estimated white blood cell proportions (using the Bakulski reference-based method^41^ for cord blood and the Houseman method^42^ for peripheral blood) including CD4+ T-lymphocytes, CD8+ T-lymphocytes, natural killer cells, B-lymphocytes, monocytes and granulocytes (nucleated red blood cells were not further analyzed due to their specificity to cord blood), gestational age, sex of the child, and cohort. Continuous covariates (estimated white blood cells, gestational age) were z-score standardized.

### Epigenetic clocks

We used several selection and exclusion criteria for the inclusion of epigenetic clocks. First, to accommodate our research objective, i.e. to characterize the developmental dynamics of sites in epigenetic clocks that have been related to age-related disease and mortality, we only included sites from epigenetic clocks that (i) have been related to age-related diseases or mortality (excluding gestational age clocks and pediatric clocks). Second, to be able to make relevant comparisons with our longitudinal DNAm set in in childhood, we (ii) only selected epigenetic clocks that have been based on human material and (iii) trained (at least in part) on DNAm extracted from blood. Furthermore, to keep our enrichment analyses balanced, we (iv) excluded heavily minimized clocks, meaning clocks designed to include a minimum number of sites and typically containing 10 or less sites (excluding Weidner’s minimized clock of three sites^12^ and Zhang’s Mortality Clock of ten sites^43^), and (v) excluded clocks that performed little feature selection (excluding Zhang’s Best Linear Unbiased Prediction clock, which includes >300,000 sites^11^). Last, we only (vi) included clocks of which the included sites are publicly available (excluding GrimAge^44^).

As a result, we selected a set of four first-generation clocks, i.e. clocks trained to estimate age: Horvath’s clock^9^ (353 sites), Hannum’s clock^10^ (71 sites), Weidner’s (non-minimized) clock^12^ (102 sites), Zhang’s Elastic Net clock^11^ (514 sites) as well as two second-generation clocks, i.e. clocks trained to estimate mortality, time-to-death, or indicators of mortality: PhenoAge^14^ (513 sites), and the DNAmTL^15^ (140 sites) and a third-generation clock DunedinPACE^16^ (173 sites), i.e. a clock trained on change estimates of age biomarkers. Since second- and third-generation clocks are both trained on (indicators of) age-related diseases or mortality, instead of age itself, these clocks were grouped together. Taken together, the first-generation clocks consisted of 967 unique sites, and the second/third-generation clocks consisted of 821 unique sites.

### Genetic and prenatal environmental factors

Methylation quantative trait locus (meQTL) associations were identified using the meqtldb database^23^, which has been generated using the ALSPAC cohort. We used (*cis*- and *trans*-) meQTLs as determined on genetic-DNAm variations in cord blood (*n*=771), as this was the tissue relevant to this analysis. As a sensitivity analysis, we also tested for (*cis*- and *trans*-) meQTLs as identified by the larger GoDMC consortium, including DNAm from cord blood and peripheral blood from participants of all ages (*n*=32,851). Site-level associations with environmental factors were identified using summary statistics^25^ from EWAS meta-analyses on prenatal environmental factors and DNAm in cord blood performed by the Pregnancy and Childhood Epigenetics (PACE) consortium^24^, that were available in the EWAS Catalog^25^ and had reported at least one association in a cell-type adjusted analysis. The EWAS Catalog includes sites with an association of *p*<1×10^−04^, yet the inclusion thresholds vary somewhat between the studies. Prenatal exposures included sustained maternal smoking (*n*=2,620 associated sites; *p*-value range=1.35×10^−206^-9.98×10^−05^)^26^, hypertensive disorders (*n*=920; *p*=1.90×10^−12^-9.90×10^−05^)^27^, overweight/obesity (*n*=159, *p*=3.80×10^−12^-1.00×10^−07^)^28^, maternal BMI (*n*=104, *p*=6.00×10^−14^-1.00×10^−07^)^28^, (pregnancy related) anxiety (*n*=57, *p*=5.33×10^−06^-9.91×10^−05^)^29^, haemoglobin levels (*n*=40, *p*=1.00×10^−07^-1.40×10^−05^)^30^, air pollution exposure, measured by NO_2_ (*n*=24, *p*=8.65×10^−08^-1.81×10^−05^)^32^, and PM_2.5_ (*n*=6, *p*=8.30×10^−08^-1.80×10^−05^)^31^.

### Enrichment analyses

Clock sites were tested for enrichment of detected (non-linear) change during development, the presence of individual variation at birth, individual variation in rate of change, meQTL associations, or associations with prenatal environmental factors. For our analysis of change, we reversed the coefficients of DNAmTL to have a consistent meaning as those of the other clocks, as telomere length decreases with age. Enrichment was determined by comparing clock sites with ‘non-clock’ sites on the 450K array with a Fisher’s exact tests. The significance threshold was set at *p*<0.05.

## Acknowledgements

The Generation R Study is conducted by Erasmus MC, University Medical Center Rotterdam in close collaboration with the School of Law and Faculty of Social Sciences of the Erasmus University Rotterdam, the Municipal Health Service Rotterdam area, Rotterdam, the Rotterdam Homecare Foundation, Rotterdam and the Stichting Trombosedienst & Artsenlaboratorium Rijnmond (STAR-MDC), Rotterdam. We gratefully acknowledge the contribution of children and parents, general practitioners, hospitals, midwives and pharmacies in Rotterdam. The generation and management of the Illumina 450K methylation array data (EWAS data) for the Generation R Study was executed by the Human Genotyping Facility of the Genetic Laboratory of the Department of Internal Medicine, Erasmus MC, the Netherlands. We thank Mr. Michael Verbiest, Ms. Mila Jhamai, Ms. Sarah Higgins, Mr. Marijn Verkerk and Dr. Lisette Stolk for their help in creating the EWAS database. We thank Dr. A. Teumer for his work on the quality control and normalization scripts.

We are extremely grateful to all the families who took part in the ALSPAC study, the midwives for their help in recruiting them, and the whole ALSPAC team, which includes interviewers, computer and laboratory technicians, clerical workers, research scientists, volunteers, managers, receptionists and nurses.

## Author contributions

R.H.M. contributed to the conception and design of the work, the analysis, interpretation of the data, drafted the work and substantively revised it. M.S. contributed to the conception and design of the work, the interpretation of the data, and substantively revised the work. A.N., and J.F.F. contributed to the interpretation of the data and substantively revised the work. C.A.M.C. contributed to the conception and design of the work, the interpretation of the data, drafted the work and substantively revised it.

All authors approve the submitted version and agree to be personally accountable for the author’s own contributions and ensure that questions related to the accuracy or integrity of any part of the work, even ones in which the author was not personally involved, are appropriately investigated, resolved, and the resolution documented in the literature.

## Data Availability

Site-level results have been made publicly available at http://epidelta.mrcieu.ac.uk/. Individual-level data from the Generation R Study are available upon reasonable request to the director of the Generation R Study, subject to local, national and European rules and regulations. ALSPAC data access is through a system of managed open access. The ALSPAC access policy (http://www.bristol.ac.uk/media-library/sites/alspac/documents/researchers/data-access/ALSPAC_Access_Policy.pdf) describes the process of accessing the data and samples in detail, and outlines the costs associated with doing so.

