## Supplementary Figure 1 for "What makes clocks tick? Characterizing developmental dynamics of adult epigenetic clock sites"

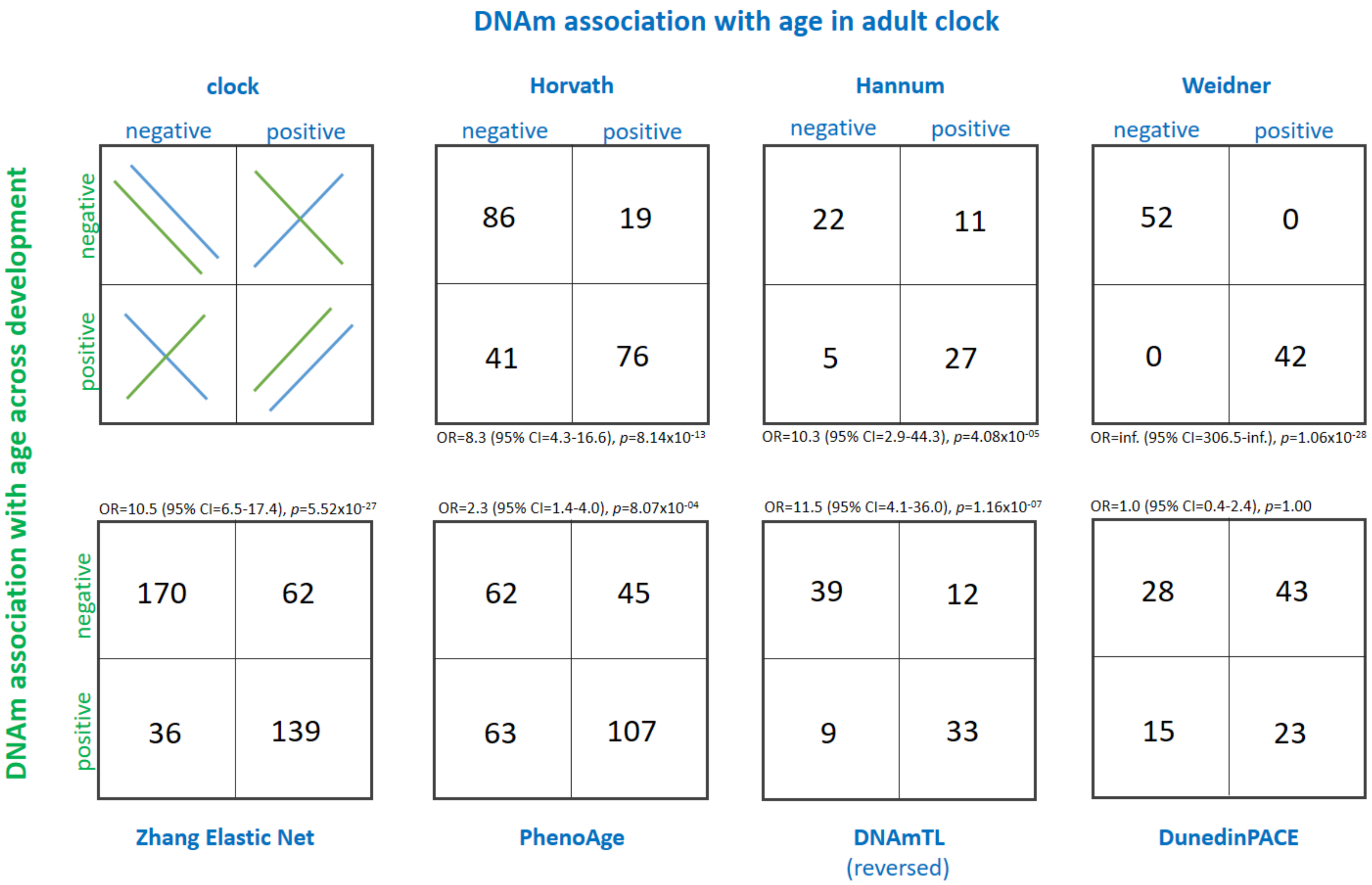


**Supplementary Figure 1.** Agreement of directionality of DNAm association with age in adult clocks and DNAm association with age across development, among sites that show significant change ($p<1x10^{-07}$) across development. Odds ratio statistics are produced using Fisher exact tests.
