## Supplementary Information for "What makes clocks tick? Characterizing developmental dynamics of adult epigenetic clock sites"

**Setting**

Epigenetic data were obtained from two prospective population-based cohorts: the Generation R Study (Generation R) in The Netherlands and the Avon Longitudinal Study of Parents and Children (ALSPAC) in the United Kingdom. For Generation R, Pregnant women residing in the study area of Rotterdam, The Netherlands, with an expected delivery date between April 2002 and January 2006 were invited to enroll. A more extensive description of the study is detailed elsewhere[1,2]. The Generation R Study is conducted in accordance with the World Medical Association Declaration of Helsinki and has been approved by the Medical Ethics Committee of Erasmus MC, University Medical Center Rotterdam. Informed consent was obtained for all participants. In the ALSPAC study, pregnant women residing in the study area of former county Avon, United Kingdom, with an expected delivery date between April 1991 and December 1992 were invited to enroll. Detailed information on the study design can be found elsewhere[3,4]. The ALSPAC website contains the details of all available data through a fully searchable data dictionary and variable search tool (http://www.bristol.ac.uk/alspac/researchers/our-data/). Ethical approval for the study was obtained from the ALSPAC Ethics and Law Committee and the Local Research Ethics Committees. Consent for biological samples has been collected in accordance with the Human Tissue Act (2004). Informed consent for the use of data collected via questionnaires and clinics was obtained from participants following the recommendations of the ALSPAC Ethics and Law Committee at the time.

In Generation R, pregnant mothers had 9,749 live-born children. Epigenetic data was available for a subsample of 1,414 children. This subsample consisted of participants with parents born in the Netherlands (European ancestry[5] confirmed for all children with genetic data available (95.4%)). Fifteen sibling pairs were present in the dataset. From each pair, one sibling with the lowest number of DNAm measurements, or otherwise randomly, was excluded, resulting in a sample with 1,399 children, including 2,333 measurements at birth and/or 6 years and/or 10 years of age.

In the ALSPAC study, the initial number of pregnancies enrolled was 14,541. Of these, 13,988 children were alive at 1 year of age. When the oldest children were approximately 7 years of age, an attempt was made to bolster the initial sample with eligible cases who had failed to join the study originally. The phases of enrolment are described in more detail elsewhere[3,4]. The total sample size for analyses using any data collected after the age of seven is therefore 15,447 pregnancies, resulting in 15,658 foetuses. Epigenetic data was available for a subsample of 1,003 children as part of the Accessible Resource for Integrated Epigenomic Studies (ARIES) study[6]. From this sample, 48 children with non-European ancestry as based on genetic principle component analysis and 6 children with missing data on gestational age were excluded, resulting in a sample of 949 children, including 2,686 measurements at birth and/or 7 years and/or 17 years. All children had European ancestry. Together, these data formed a single dataset consisting of 2,348 children with 5,019 measurements.
