## Supplementary Table 1 for "What makes clocks tick? Characterizing developmental dynamics of adult epigenetic clock sites"

**Supplementary Table 1. Enrichment patterns for epigenetic clocks**

| | | First-generation clocks | | | | | | | | | | | | Second-generation clocks | | | | | | Third-generation clock | | |
|---|---|---|---|---|---|---|---|---|---|---|---|---|---|---|---|---|---|---|---|---|---|---|
| | All autosomal sites o | Horvath's clock sites (*n* =353)[1] | | | Hannum's clock sites (n=71)[2] | | | Weidner's clock sites (n=102)[3] | | | Zhang's clock sites (n=514)[4] | | | PhenoAge clock sites (n=513)[5] | | | Telomere clock sites (n=140)[6] | | | DunedinPACE clock sites (n=173)[7] | | |
| | % yes | % yes | OR | *p* -value | % yes | OR | *p* -value | % yes | OR | *p* -value | % yes | OR | *p* -value | % yes | OR | *p* -value | % yes | OR | *p* -value | % yes | OR | *p* -value |
| DNAm change between birth and early adulthood | 51.55 | 62.89 | 1.59 | 1.91E-05 | 91.55 | 10.18 | 5.06E-13 | 98.99 | 92.14 | 3.41E-27 | 79.18 | 3.58 | 2.47E-38 | 54.19 | 1.11 | 2.33E-01 | 68.38 | 2.03 | 9.81E-05 | 63.01 | 1.60 | 2.90E-03 |
| Non-linear DNAm change - change until age 6 | 7.95 | 13.60 | 1.82 | 3.43E-04 | 14.08 | 1.90 | 7.37E-02 | 43.43 | 8.90 | 1.20E-21 | 26.85 | 4.26 | 1.71E-37 | 12.28 | 1.62 | 7.54E-04 | 27.21 | 4.33 | 2.04E-11 | 26.59 | 4.20 | 2.00E-13 |
| Non-linear DNAm change - at age 9 | 2.67 | 7.08 | 2.78 | 1.27E-05 | 8.45 | 3.37 | 1.18E-02 | 9.09 | 3.65 | 1.37E-03 | 9.53 | 3.85 | 3.60E-14 | 4.87 | 1.87 | 5.40E-03 | 2.21 | 0.82 | 1.00E+00 | 5.78 | 2.24 | 2.77E-02 |
| Inter-individual differences in DNAm level at birth | 26.21 | 33.14 | 1.40 | 3.63E-03 | 33.80 | 1.44 | 1.76E-01 | 48.48 | 2.65 | 2.85E-06 | 50.39 | 2.86 | 1.91E-31 | 38.60 | 1.77 | 9.28E-10 | 72.06 | 7.26 | 8.26E-29 | 83.82 | 14.59 | 1.46E-56 |
| Inter-individual differences in rate of DNAm change from birth | 3.35 | 4.25 | 1.28 | 3.71E-01 | 16.90 | 5.87 | 4.02E-06 | 18.18 | 6.42 | 5.19E-09 | 14.40 | 4.87 | 1.19E-25 | 4.29 | 1.29 | 2.20E-01 | 13.97 | 4.69 | 1.64E-07 | 7.51 | 2.35 | 8.55E-03 |
| Inter-individual differences in rate of DNAm change from age 6 | 0.17 | 0.00 | 0.00 | 1.00E+00 | 0.00 | 0.00 | 1.00E+00 | 0.00 | 0.00 | 1.00E+00 | 0.19 | 1.16 | 5.78E-01 | 0.39 | 2.33 | 2.13E-01 | 0.74 | 4.41 | 2.04E-01 | 0.00 | 0.00 | 1.00E+00 |
| Inter-individual differences in rate of DNAm change from age 9 | 8.17 | 6.80 | 0.82 | 3.83E-01 | 28.17 | 4.41 | 6.44E-07 | 7.07 | 0.86 | 8.54E-01 | 9.53 | 1.19 | 2.59E-01 | 7.21 | 0.87 | 4.68E-01 | 6.62 | 0.80 | 6.38E-01 | 1.16 | 0.13 | 1.24E-04 |
| meQTL associations at birth | 7.93 | 12.75 | 1.70 | 1.54E-03 | 2.82 | 0.34 | 1.25E-01 | 12.12 | 1.60 | 1.33E-01 | 12.26 | 1.62 | 5.77E-04 | 12.87 | 1.72 | 1.10E-04 | 27.94 | 4.51 | 4.16E-12 | 28.90 | 4.73 | 4.29E-16 |
| Prenatal environmental exposure associations | 0.81 | 1.70 | 2.11 | 7.03E-02 | 1.41 | 1.74 | 4.40E-01 | 5.05 | 6.50 | 1.34E-03 | 3.70 | 4.70 | 7.85E-08 | 0.97 | 1.20 | 6.17E-01 | 3.68 | 4.66 | 5.29E-03 | 4.62 | 5.93 | 9.74E-05 |
